# Regional dynamics in the evapotranspiration components, crop coefficients and water productivity in vineyards in the Barossa Valley

**DOI:** 10.1101/2024.06.09.598151

**Authors:** V. Phogat, P.R. Petrie, M. Bonada, C. Collins

**Affiliations:** Crop Sciences, South Australian Research and Development Institute, GPO Box 397, Adelaide, SA 5001, Australia; School of Agriculture, Food and Wine, Waite Research Institute, University of Adelaide, Waite Campus, PMB1 Glen Osmond, SA 5064, Australia; College of Science and Engineering, Flinders University, Adelaide SA 5000, Australia; ARC Training Centre for Innovative Wine Production, University of Adelaide, PMB1 Glen Osmond, SA 5064, Australia

**Keywords:** Grapevine, irrigation, FAO-56, actual evapotranspiration, transpiration, crop coefficients, water stress coefficient, water productivity

## Abstract

Estimation of water balance components, water stress and crop coefficients at different spatial scale are crucial for understanding regional dynamics in irrigation requirement and crop water use. We estimated these parameters for irrigated vineyards over 3 consecutive seasons (2018-19, 2019-20 and 2020-21) at 48 locations in the Barossa region, South Australia. We used FAO-56 dual crop coefficient approach by integrating relevant data for soil, crop, and climate parameters from the study sites. Numerous statistical error estimates, and efficiency parameters were estimated to compare and verify the predictions by FAO-56 approach. Results show a huge variability in the irrigation, water balance parameters, crop and water stress coefficients, and water productivity parameters. For instance, a coefficient of variation ranging from 20 to 97% was observed in daily and seasonal actual ET (*ET*_*c act*_) across different sites and seasons. Average actual transpiration (*T*_*p act*_) and evaporation (*E*_*s*_) account for around 65 and 35% of the *ET*_*c act*_, respectively, showing the potential to save water lost to the environment from the soil surface. Estimated actual single crop coefficient (*K*_*c act*_) across all sites varied from 0.35 to 0.59, 0.16-0.62 and 0.18-0.68 during the budburst to flowering (BB-FL), flowering to veraison (FL-V), and veraison to harvest (V-H) stages of crop growth, respectively. Similarly, actual basal crop coefficients (*K*_*cb act*_) for grapevine reveal immense site-specific variability questioning the adoption of uniform coefficients at subregional and regional levels. Results further demonstrate that water stress (*K*_*s*_) gradually increased reaching its peak from late November to early December, with variations across the region ranging from 23 to 64%. A comparison of water productivities in relation to *ET*_*c act*_ and *T*_*p act*_ exhibit almost 61% higher values for the latter across all the sites and subregions. Dry biomass productivity shows huge potential for renewal energy generation. Variations in the components of ET and crop coefficients are consistent with the characteristic variation in soil, topography, and microclimates. This study suggests that locally estimated *K*_*c act*_ and *K*_*cb act*_ will contribute to the efficient use of limited freshwater resources for sustainable wine grape production.

## 1. Introduction

Grapevine (*Vitis vinifera* L.) is one of the most important horticultural crops which is grown in diverse climate across different continents (Sadras, 2012) covering approximately 7.20 million ha area (Organization of Vine and Wine, 2023). Many of these vineyards are located in arid, semiarid, and Mediterranean agroecosystems where evapotranspiration is a major component of the water cycle. However, most of the grape growing regions are facing enormous challenges for adequate fresh water under limiting climatic conditions and shortfall in the seasonal precipitation leading to deficit water condition in the soil (Romero et al., 2022). Lack of optimal soil moisture regime is described as one of the most important constraints limiting water uptake, growth, maturity, production, and fruit and wine quality (Castellarin et al., 2007; Ramos et al., 2020; Roby et al., 2004; Yang et al., 2022).

In Australia, water availability for irrigation is highly erratic and uncertain. Seasonal water allocation to irrigated industries from the Murray Darling River system depends on the rainfall and water flow conditions. For example, during Millennium drought (2002-2009) the water allocation was severely reduced, to a level as low as 18% of normal allocations. Uncertain water allocations can have severe impact on the sustainable production and resilience of irrigated crop including vineyards in the Barossa Valley region which imports 70 % of irrigation water from the Murray River system (Department of Environment and Water, 2022). Climatic conditions are expected to deteriorate further due to variability in the Indian Ocean Dipole and El Nino Southern Oscillation which are the key drivers of Australian climate (e.g., (King et al., 2020). Several studies have concluded that climate induced water scarcity is expected to have unprecedented impact on water availability, yield, and wine quality (IPCC, 2022; Jones et al., 2022; Phogat et al., 2018; Santos et al., 2020; Webb et al., 2012). It is well understood that grape’s productivity strictly depends on water availability which is fundamental for vine’s growth and development of berries with chemical and physical features assuring high wine quality (Chacón-Vozmediano et al., 2020). Hence, site-specific accurate estimation of evapotranspiration (*ET*) and its partitioning into transpiration (*T*_*p*_) and evaporation (*E*_*s*_) and driving location specific crop coefficients (K_c_) is fundamental for improving water management practices in the water-limited environments and under deficit irrigation conditions.

A wide range of methods and techniques have been used for estimating water relations, ET components and crop coefficients of grapevine which include lysimeter (Hochberg et al., 2023; Munitz et al., 2019; Netzer et al., 2009; Poblete-Echeverría and Ortega-Farias, 2013; Zhang et al., 2011), sap flow (Ferreira et al., 2012; Intrigliolo and Castel, 2009; Wang et al., 2019; Yunusa et al., 2004), Bowen ratio (Zhang et al., 2011), surface energy balance (Castellví and Snyder, 2010; Moratiel and Martínez-cob, 2012), eddy covariance (Ferreira et al., 2012; Ortega-Farias et al., 2010; Poblete-Echeverría and Ortega-Farias, 2013; Wang et al., 2019), soil water balance (e.g., (Fooladmand and Sepaskhah, 2009), remote sensing and unmanned aerial vehicle (UAV) involving image analysis (Beeri et al., 2020; Carrasco-Benavides et al., 2012; Matese et al., 2022; Poblete-Echeverría et al., 2017; Pôças et al., 2020; Romero et al., 2018), and a combination of methods (Cancela et al., 2012; Poblete-Echeverría and Ortega-Farias, 2013; Trambouze et al., 1998). These methods have their own inherent pros and cons when used for the estimation of water balance, soil water regimes, and dynamics of water deficit conditions in the soil at varied spatial scale (Pereira et al., 2021c; Pôças et al., 2020). In fact, intensive field measurements of water balance components require large investments in sensors, labour, and time which is prohibitive to adopt at regional scale keeping in view the large-scale heterogeneities in soil and variabilities in growing conditions. Moreover, climate variability, canopy characteristics and heterogeneity in soil characteristics varying at spatial scale play a critical role in estimating accurate crop irrigation requirement and understanding the extent of variability in crop water demand (Fraga et al., 2017; Zhu et al., 2021). Therefore, a combination of site-specific field measurements and simple modelling tools can provide useful information for understanding the inherent dynamics of water balance components and improving the efficient use of scarce water resources.

There are numerous modelling approaches to estimate the plant water uptake and water balance components under cropped conditions used as an alternative to the complex field measurement methods. Among them, FAO-56 (Allen, 1998) has been extensively used to derive crop evapotranspiration (*ET*_*C*_), crop transpiration (*T*_*p*_), soil evaporation (*E*_*s*_), irrigation requirement and local coefficients of different crops under varied water availability conditions due to its simplicity and ease of application (Cancela et al., 2015; Fandiño et al., 2012; Phogat et al., 2020a; Wang et al., 2019). This method is accepted as a standard technique (Pereira et al., 2015) over the complex field methods and has been used extensively to calculate *ET*_*C*_ demand, ET partitioning, *E*_*s*_ losses and irrigation schedule of the entire range of crops (Pereira et al., 2021b; Pereira et al., 2021c; Rallo et al., 2021) including grapevines (Fandiño et al., 2012; Phogat et al., 2020a; Phogat et al., 2017; Picón-Toro et al., 2012; Poblete-Echeverría and Ortega-Farias, 2013; Zhao et al., 2018), and other horticultural crops such as apple ((Marsal et al., 2012; Mobe et al., 2020; Zanotelli et al., 2019), almonds (e.g., (Drechsler et al., 2022; Phogat et al., 2013), citrus (Er-Raki et al., 2009), peach (Paço et al., 2011), and olives (Conceição et al., 2017; Er-Raki et al., 2010; Paço et al., 2019; Paço et al., 2014). This technique is also employed to estimate potential *T*_*p*_ and *E*_*s*_ which serves as crucial inputs in agro-hydrological modelling studies (Kool et al., 2014; Phogat et al., 2023; Phogat et al., 2017; Ramos et al., 2012) and for assessing the climate change risks to irrigated viticulture (Phogat et al., 2018; Phogat et al., 2020b). Good agreement has been reported by several studies between FAO-56 and other methods based on field measurements of water consumption by multiple crops (Pereira et al., 2021a; Pereira et al., 2021b; Rallo et al., 2021). Improvements in FAO-56 approach in estimating crop evapotranspiration (*ET*_*C*_) and related parameters are explained in subsequent publications (Allen and Pereira, 2009; Allen, 2005; Allen et al., 2005; Pereira et al., 2020; Phogat et al., 2016). Typically, the values of crop coefficients (K_c_ and K_cb_) are reported to be sensitive to climate, crop management, soil type, hydrological and environmental factors (e.g., (Pereira et al., 2021b; Rallo et al., 2021).

However, estimating *ET*_*C*_ and irrigation requirement of crops using the FAO–56 proposed generic *K*_*c*_ and *K*_*cb*_ values may induce significant uncertainties in crop water use (Taylor et al., 2014; Widmoser, 2009; Yang et al., 2016) and therefore, integration of estimated crop coefficients adapted to local soil, crop and climate conditions is essential to obtain reliable information (Pereira et al., 2021b; Rallo et al., 2021). Use of location specific basal (*K*_*cb*_) and single (*K*_*c*_) crop coefficients values for irrigation scheduling helps in the precise estimation of the dynamics of evaporation (*E*_*s*_) and transpiration (*T*_*p*_) leading to judicious planning and management of scarce water resources in drip-irrigated high-value cropping systems (Carrasco-Benavides et al., 2012; Phogat et al., 2017). There can be an immense variability in the site-specific vine water requirement within a region and losses of water through *E*_*s*_, particularly in sparsely planted trees and orchards, such as grapevines, where a large soil surface area is exposed to atmospheric effects. Besides *E*_*s*_, *T*_*p*_, *K*_*c*_ and *K*_*cb*_ a reliable understanding of site-specific water stress coefficient (*K*_*s*_) is critical for providing real time irrigation schedule under reduced water allocation across the region, especially under deficit conditions being practiced in the vineyards for better wine quality. Therefore, reliable estimation of these components at regional scale can help providing useful information for irrigation management, improving water use efficiency and sustainable management of scarce water resources.

The objectives of this study, thus, were (i) to calibrate and validate FAO-56 DCC model for dynamics profile water availability under grapevines at multiple locations (24) in the Barossa region and (ii) to estimate actual *ET*_*C*_, *T*_*p*_, *E*_*s*_, seasonal water stress (K_s_), and location specific crop coefficients (*K*_*c*_ and *K*_*cb*_) of grapevine in the Barossa region over multiple seasons (2018-19, 2019-20, 2020-21) and locations (24), and (iii) to compare the water balance components, crop coefficients and imposed stress level across the Barossa region to understand the inherent variability induced by soil type, vine canopy and climate parameters. This information would help improve the irrigation scheduling and better utilization of water resources for irrigation in the Barossa region.

## Materials and Methods

### Description of study region

The study area (Barossa region) is recognised as a leading wine grape producing region in Australia. The region experiences a Mediterranean climate, characterised by hot, dry summers and cool to cold winters. Long-term (124 years) average rainfall in the region is 467 mm at Bureau of Meteorology station 023343 within the Barossa region (https://www.bom.gov.au/jsp/ncc/cdio/cvg/av) and annual evapotranspiration is 1300 mm, resulting in the need for irrigation for crop production. Most of the rainfall is occurring in the winter months coinciding with the dormant stage of grapevine. During the summer months grapevines requires supplemental irrigation which coincides with the typical canopy growth and growing season climate.

Across the Barossa GI Zone (which includes Barossa and Eden Valley regions), viticultural management practices including irrigation amounts vary immensely for achieving a characteristic vine style, yield components and wine quality attributes influenced by the individual climate, soil, topography, genotype and management conditions. Varied levels and timing of water stress is applied by the growers which significantly impact the site-specific irrigation requirement and growing conditions. A large-scale heterogeneity in the soil across the region with varied extent of plant available water capacities also play an important role in adopting a typical irrigation schedule for establishing a grapevine orchard (Phogat et al., 2024).

### Soil sampling and analysis

Twenty-four sites within the grapevine orchards were identified with varied growing conditions which broadly covers the diverse climate-soil-crop management practices implemented by the growers (Collins et al., 2022). Twenty sites were selected from the various sub-regions within the Barossa Valley (Robinson and Sandercock, 2014) while four sites (21 to 24) were spread within the Eden valley region (Fig. 1). Both regions are the part of broader Barossa valley GI zone (Wine Australia, 1996). At each of the study site, two distinctively different grapevine orchard sites were selected to integrate local soil and canopy variability within the orchards. Soil profile sampling was carried out during June to August 2019 at each of the study site, totalling 48 profile samples. The samples were collected using a 4 cm internal diameter hydraulic hammer-driven probe. Cores were collected from vineyards at 0.4 m from the drip irrigation line in the mid-row and between the emitters. Two intact cores up to 90 cm depth were taken in PVC clear sleeves from each vineyard soil profile site using a Geoprobe Macro-Core® soil sampling system (Earthtech Drilling Products Pty. Ltd., Victoria, Australia). At some locations, sampling was restricted to shallower depths due to the presence of rocks. The cores were detached from the assembly and capped immediately before their transfer to the laboratory. Core samples were oven dried at 40°C, caped and stored at room temperature until processing.

**Fig. 1.**
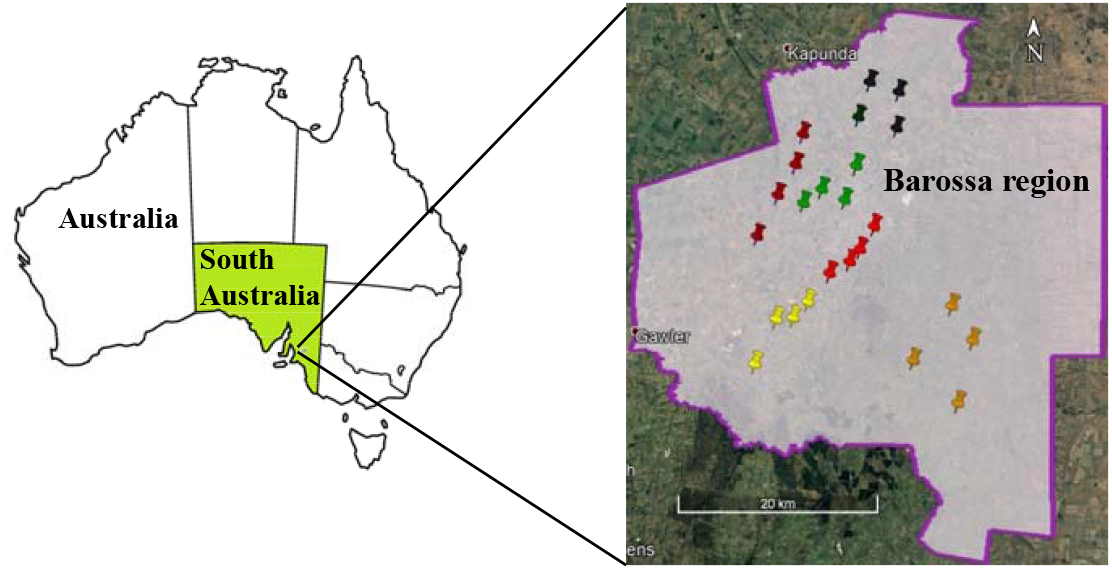
Schematic map showing study sites in the Barossa region. Study sites are colour coded to represent different sub-regions: northern grounds (grey), central grounds (green), eastern edge (red), southern grounds (yellow), western ridge (purple) and Eden Valley (orange).

Soil sub-samples were drawn from the cores based on visual variation in texture and distinct horizons for determining the physico-chemical characteristics of the soils. For bulk density determination, clods were collected from the plastic cores in duplicate and stored in the moisture boxes for further analysis. Particle size analysis was performed for each of the soil sample following the MIR spectroscopy and least square analysis (Janik et al., 2016). Bulk density was determined following the clod method (Cresswell and Hamilton, 2002). Other soil properties (OC, CEC, Exchangeable Na, Ca, Mg, K, ESP, B) were estimated following the standard procedures outlined in (Rayment et al., 2011).

Details of soil phyco-chemical characterisation including water content-pressure head relationship and the estimation of plant available water capacity (PAWC) following fixed and dynamics field capacity methods for all the study sites are described in detail in (Phogat et al., 2024). Daily plant available water (PAW) in the soil profile was estimated by subtracting the estimated depth of water at CLL for each texture layer from the corresponding total available water (TAW) estimated from the probe data. Plant available water showed strong seasonal response in relation to the rainfall, irrigation, and seasonal evapotranspiration fluxes during the wine grape season. It is worth noting that estimated diverse site-specific water retention properties of the soil may have remarkable influence on the water availability to wine grapes in the Barossa region (Phogat et al., 2024).

### Monitoring soil water content dynamics

Soil moisture probes (EnviroSCAN, Sentek technologies) were installed at all the 48 sites for monitoring the real-time water content dynamics in the soil. These probes were fitted with 4 sensors positioned at 20, 50, 60-80 and 70-110 cm depths depending on profile depth at different locations. Site-specific calibration was performed for all the probes. Volumetric soil moisture was monitored and logged at daily intervals recording changes in soil moisture at different depths. This allows to estimate the real-time water content distribution in the soil and the extent of daily soil moisture deficit in the soil profile. The daily values of profile plant available water (PAW) were estimated from the changes in water content data. Daily PAW served as observed values for comparison with corresponding values estimated by the FAO-56 DCC approach (Allen, 1998) adopted in this study for partitioning the crop evapotranspiration and the estimation of site-specific crop coefficients under local climate and growing conditions.

### Climate data

Portable automatic weather stations were installed at each site (MEA Junior WS, Magill, South Australia) which measured the daily values of temperature, relative humidity, rainfall, wind speed and solar radiation. Reference crop evapotranspiration (*ET*_*0*_) was estimated following Modified Penman and Monteith outlined in Allen et al. (1998). Seasonal (15^th^ September to 31^st^ March) values of rainfall and *ET*_*0*_ measured during the 2018-19, 2019-20, and 2020-21 at each location are shown in Fig. 2.

**Fig. 2.**
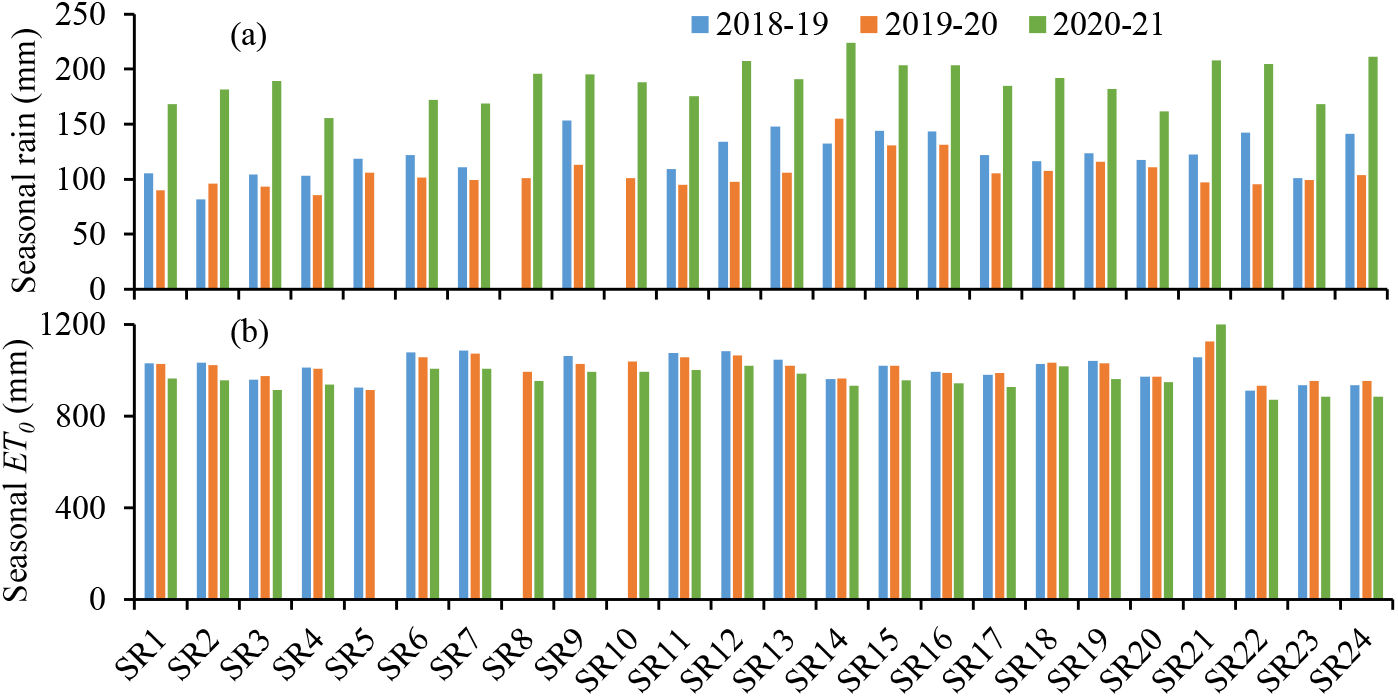
Seasonal (15^th^ September to 31^st^ March) values of (a) rainfall and (b) reference crop evapotranspiration (ET_0_) during the 2018-19, 2019-20 and 2020-21 at different sites in the Barossa region.

Seasonal rainfall (15^th^ September-31^st^ March) across the study sites during 2018-19 and 2019-20 was almost similar while the values increased by 65% during 2020-21 as compared to the average of previous two years. However, site specific variability existed, and values were varied from 81.8 to 144, 86-131, 156-224 mm during the 2018-19, 2019-20 and 2020-21, respectively. Spatial variation in the average rainfall existed from north-east (sites 1-4) to south-west (sites 13-16) direction showing increasing trend over the three seasons. Seasonal *ET*_*0*_ values during the 2018-19, 2019-20 and 2020-21 seasons were 1010.2, 1010.2 and 968.9 mm, respectively. These values varied in a narrow range across the study sites except SR21 where slightly higher values of *ET*_*0*_ were recorded compared to other sites during all the three seasons.

### Canopy measurements

Seasonal canopy growth of grapevines was monitored on fortnightly basis at all the locations and leaf area index (LAI) was estimated from bud burst till the harvest of berries. Images of the canopy were taken by iPhone from beneath the vines looking upward during the early morning to avoid overexposure of the image sections by the sun. These images were analyzed using the ImageJ software (https://imagej.nih.gov/ij/index.html version 1.53k/Java1.8.0_172; accessed 27 December 2023). The canopy size for the entire growing season was indirectly assessed as the leaf area index (*LAI*), following the methodology described in (De Bei et al., 2016; Fuentes et al., 2014; Phogat et al., 2023).

### FAO-56 Dual crop coefficient approach

FAO-56 outlines two approaches, the single and the dual crop coefficients (FAO-56 DCC) approach. The FAO-56 DCC method is particularly suitable for crops having incomplete ground cover or where a fraction of the soil surface is wetted by the irrigation and exposed to radiation (Allen, 2005), such as under drip irrigated grapevines. This approach estimates plant transpiration and soil evaporation separately involving separate coefficients for transpiration (basal crop coefficient, *K*_*cb*_) and evaporation (soil evaporation coefficient, *K*_*e*_). The FAO-56 DCC methodology requires a considerable amount of data for crop, soil, and climate parameters. Detailed procedure is outlined in Allen et al. (1998) and subsequent improvements are described in other publications (Allen and Pereira, 2009; Allen, 2005; Pereira et al., 2021c).

The *ET*_*C*_ in FAO-56 DCC approach is estimated as:

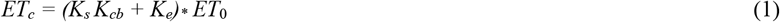

The generic values of *K*_*cb*_ (dimensionless) reported in FAO-56 needs to be modified for the climate and crop canopy cover, height and density of vegetation and amount of stomatal regulation under wet soil condition (Allen and Pereira, 2009; Pereira et al., 2020). Under reduced water availability in the soil *K*_*cb*_ is further modified in accordance with stress reduction coefficient (*K*_*s*_) estimated from profile water balance. On the other hand, evaporation component (*K*_*e*_) from the soil surface depends on the dynamics fraction of wetted and exposed soil surface which is related to the vegetation cover and extent of rainfall and irrigation applied to the crop.

The basal crop coefficients for the agricultural crop can be adjusted to the local climate conditions using the following equation:

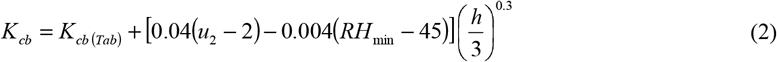

where, *K*_*cb (Tab)*_ is the tabulated value of *K*_*cb mid*_ or *K*_*cb end*_ (Table 1), *u*_*2*_ is the mean value of daily wind speed during the mid or late season growth stage (m s^-1^), *RH*_*min*_ is the mean value of daily minimum relative humidity during the mid- or late season growth stage (%), and *h* is the mean plant height during the mid or late season growth stage (m).

The basal *K*_*cb*_ represents transpiration water loss and is correlated with the amount of vegetation and can be expressed in terms of a crop density coefficient, *K*_*d*_ (Allen et al., 2007; Allen and Pereira, 2009):

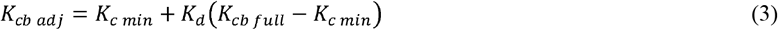

where *K*_*cb full*_ is the estimated basal *K*_*cb*_ (eq. 3) under peak plant growth conditions (LAI>1) and local climate conditions, and *K*_*c* min_ is the minimum *K*_*c*_ for bare soil whose value is 0.15 under typical agricultural conditions (Allen et al., 1998). However, for large stand size (>500 m^2^) tall vegetation having full ground cover (LAI>3) under full water supply, *K*_*cb full*_ can be approximated with plant height and adjusted for climate as below:

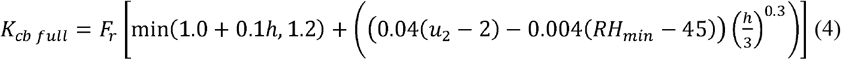

where *F*_*r*_ [0–1] is an adjustment factor relative to crop stomatal control. Basically, *F*_*r*_ applies a downward adjustment (*F*_*r*_<1.0) if the vegetation exhibits more stomatal control on transpiration than is typical of most annual agricultural crops (Allen and Pereira, 2009). It can be estimated based on the FAO Penman–Monteith equation and assuming full cover conditions (Allen et al., 1998). However, for this study, the values of *F*_*r*_ for grape vine were taken from Allen and Pereira (2009).

The *K*_*d*_ allows adjusting *K*_*cb*_ to the vegetation and ground cover conditions (Allen and Pereira, 2009) which can be estimated as:

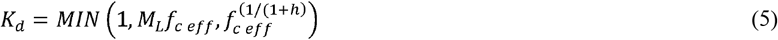

where *f*_c,eff_ is the effective fraction of ground covered shaded by vegetation near solar noon (dimensionless), *h* is crop height (m) and *M*_*L*_ is a multiplier for *f*_c,eff_, representing the ratio of *ET*_*c*_ per unit of horizontal vegetation surface to *ET*_*0*_ over the same surface (1.5-2.0). The *K*_*d*_ parameter can also be approximated under normal conditions as a function of the measured or estimated leaf area index (LAI) during mid-season (Allen et al., 1998) as:

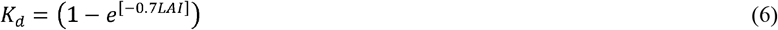

The LAI values for wine grape were estimated periodically during the growing season at each of the study sites (24) in the Barossa for the three seasons (2018-19 to 2020-21). The site-specific climate adjusted *K*_*cb*_ (*K*_*cb adj*_) values were further improved by imposing a stress reduction coefficient (*K*_*s*_). The latter is calculated from the estimated daily water balance for different sites involving initial water content in the profile, total available water (TAW) in the soil profile, readily available water (RAW) and root zone depletion (*D*_*r*_) as:

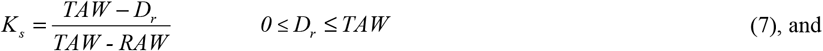

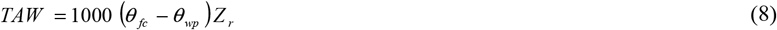

where *θ*_*fc*_ is the water content at field capacity (cm^3^/cm^3^), *θ*_*wp*_ represents the water content at wilting point (cm^3^/cm^3^) and *Z*_*r*_ is the rooting depth (m). The RAW in FAO-56 DCC is estimated as the fraction of TAW which a crop can extract from the root zone without suffering water stress, given as:

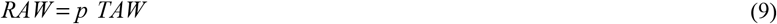

where *p* is the average fraction of TAW that can be depleted from the root zone before water stress occurs (0-1). It was estimated from a numerical approximation defined in FAO 56 as:

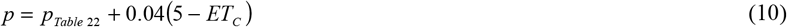

where *p*_*Table 22*_ is the tabulated value of this fraction for different crops given in FAO 56 (Table 2). Root zone depletion (*D*_*r*_) in the soil in FAO-56 method is computed through water balance at the end of every day which is given as:

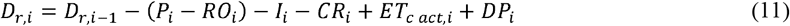

where *D*_*r,i*_ is the root zone depletion at the end of day *i* (mm), *D*_*r,i-1*_ is the root zone depletion at the end of the previous day *i*-1 (mm), *P*_*i*_ is the daily precipitation (mm) on day *i, RO*_*i*_ represents the run off (mm) of which the impact can be considered as negligible as explained in FAO-56 (Allen et al., 1998), *I*_*i*_ is the daily irrigation (mm), *CR*_*i*_ represents the capillary rise from the groundwater table on day *i* (mm) which was assumed negligible for deep water table conditions (i.e. > 2 m in this study), *ET*_*c act,i*_ is the actual crop evapotranspiration on day *i* (mm), and *DP*_*i*_ is the water loss out of the root zone by deep percolation on day *i* (mm). It is assumed that as long as the soil water content in the root zone is below field capacity, the soil will not drain and *DP*_*i*_ = 0. However, following heavy rain or excessive irrigation, *DP*_*i*_>0 and drainage takes place. The *ET*_*c act,i*_ refers to both optimal and sub-optimal crop irrigation conditions, i.e. under full or deficit irrigation.

The soil evaporation coefficient (*K*_*e*_) expresses the evaporation component of *ET*_*C*_. It attains a maximum value following precipitation or full cover irrigation, e.g. sprinkler irrigation, and becomes minimal or zero when the soil surface is dry. The crop coefficient (*K*_*c*_ *= K*_*cb*_ *+ K*_*e*_) can never exceed the maximum value (*K*_*c max*_) which is estimated from the relative humidity (*RH*_min_), wind speed (*u*_2_) and crop height (*h*) as:

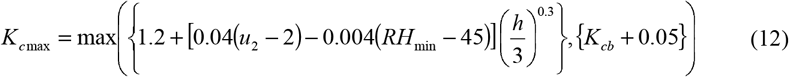

*K*_*c max*_ represents an upper limit on evaporation and transpiration from the cropped surface and reflects the natural constraints on available energy. It ranges from about 1.05 to 1.3 when using the grass reference *ET*_*0*_ (Allen et al., 2005).

Evaporation component of *K*_*c*_ (*K*_*e*_) is reduced below the potential level, due to the reduced availability of water for evaporation from the soil surface. Under these conditions *K*_*e*_ is estimated as:

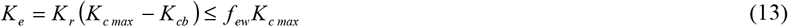

where, *K*_*r*_ is the soil evaporation reduction coefficient ranging from 0 to 1. When the soil surface is very wet as happens during precipitation events, *K*_*r*_ = 1 (stage 1 evaporation: (Ritchie, 1972)). However, a few days after a precipitation event, evaporation is reduced below the potential rate (*K*_*c max*_ *–K*_*cb*_), which represents stage 2 of evaporation (Ritchie, 1972). The parameter *f*_*ew*_ represents the exposed and wetted soil surface fraction. Where the complete soil surface is wetted, as by precipitation or sprinkler irrigation, the fraction of soil surface from which most evaporation occurs, *f*_*ew*_, is essentially defined as (1-*f*_*c*_), where *f*_*c*_ is the average fraction of soil surface covered by vegetation and (1-*f*_*c*_) is the approximate fraction of soil surface that is exposed.

For irrigation systems where only a fraction of the ground surface is wetted, *f*_*ew*_ must be limited to *f*_*w*_, the fraction of the soil surface wetted by irrigation. Therefore, *f*_*ew*_ is calculated as:

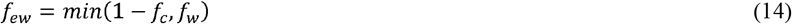

The soil evaporation reduction coefficient, *K*_*r*_ is estimated as:

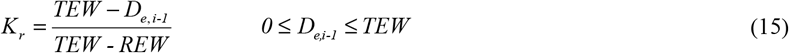

where TEW is the total evaporable water (mm) and REW represents readily evaporable water (mm). The *D*_*e,i-1*_ in Eq 15 is the cumulative depth of evaporation from the soil surface layer at the end of day *i*-1 (mm), which is estimated by the daily water balance described in FAO 56 (Eq 77 in Allen et al., 1998). The TEW is estimated as:

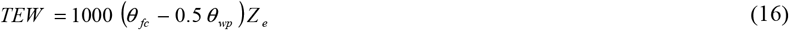

Where *θ*_*fc*_ is water content at field capacity, *θ*_*wp*_ represents water content at wilting point and *Z*_*e*_ is the evaporating depth, or the soil depth from which evaporation draws water, taken as 0.15 m (Allen et al., 1998). The TEW and REW were estimated for the surface soil layer (0.15 m) at each of the study site.

The daily water balance in *Z*_*e*_ is estimated as:

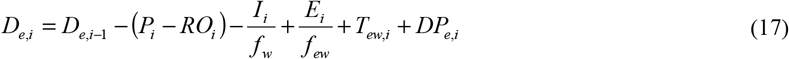

where, *D*_*e,i-1*_is the cumulative depth of evaporation following complete wetting from the exposed and wetted fraction of the topsoil at the end of day i-1 (mm), *D*_*e,i*_ is the cumulative depth of evaporation (depletion) following complete wetting at the end of day *i* (mm), *P*_*i*_ is the precipitation on day *i* (mm), *RO*_*i*_ represents precipitation runoff from the soil surface on day *i* (mm) and the impact of this fraction can be considered as negligible as explained in FAO 56 (Allen et al., 1998), *I*_*i*_ is the irrigation depth on day *i* that infiltrates the soil (mm), *E*_*i*_ represents evaporation on day *i* (i.e., *E*_*i*_ *= K*_*e*_ *ET*_*o*_) (mm), *T*_*ew,i*_ is the depth of transpiration from the exposed and wetted fraction of the soil surface layer on day *i* (mm) which is assumed as negligible for deep rooted crops (Allen et al., 1998), *DP*_*e, i*_ is the deep percolation loss from the topsoil layer (15 cm) on day *i* if soil water content exceeds field capacity (mm).

Drip irrigation applies water to only a small fraction of the soil surface, hence the calculation of the evaporation coefficient (*K*_*e*_) for drip irrigation (both surface and subsurface) requires calculation of the daily water balance for the fraction of soil surface which is wetted and exposed to sunlight (*f*_*ew*_). (Phogat et al., 2016) have calibrated and validated these fractions (*f*_*w*_, *f*_*ew*_) for wine grape for different drip line spacings.

Estimation of daily depletion (eq 11) involves the simultaneous estimation of daily crop evapotranspiration (*ET*_*C adj*_) and daily soil evaporation (*E*_*s*_) as described above. Daily soil water depletion thus obtained at each study site allows the estimation of daily residual plant available water (PAW) in the crop root zone which was calibrated against the estimated plant available water from the soil moisture probes data at each of the study sites.

### Statistical analysis

#### Error estimates

These values are compared with the model estimated values to simulate the water balance and stress level in the soil under wine grape across Barossa region. The verification of modelled values was done by estimating 3 statistical parameters i.e. mean error (*ME*), mean absolute errors (*MAE*) and root mean square error (*RMSE*). These errors were calculated from the measured and simulated daily PAW in the soil during the three wine grape seasons (2018-19, 2019-20, and 2020-21). The following equations were employed to estimate *ME, MAE* and *RMSE* by comparing measured (*M*) and corresponding model simulated (*S*) values:

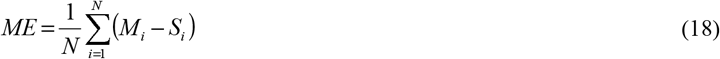

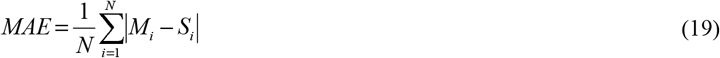

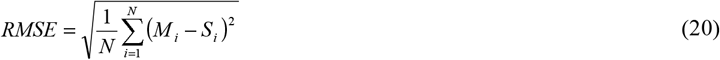

Here, *N* is the number of comparisons.

#### Efficiency criteria

The coefficient of determination (*R*^*2*^) was applied for testing the proportion of variance in the measured data explained by the model, and is defined as the square of the coefficient of correlation (*r*) according to Bravais-Pearson, calculated as:

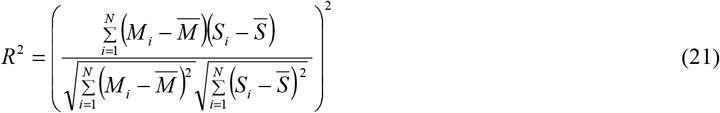

Values of *R*^*2*^ can vary between 0 and 1, with higher values indicating less variance, and values greater than 0.5 typically considered acceptable (Van Liew et al. 2003).

Model efficiency (*E*), as proposed by Nash and Sutcliffe (1970), is defined as one minus the sum of absolute squared differences between simulated and measured values, normalized by the variance of measured values during the period under investigation:

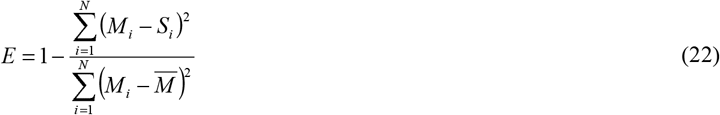

The range of *E* lies between − ∞ and 1.0 (perfect fit). An efficiency value between 0 and 1 is generally viewed as an acceptable level of performance. Efficiency lower than zero indicates that the mean value of the observed time series would be a better predictor than the model and denotes unacceptable performance (Moriasi et al. 2007). As the infinite value of E creates complexity in the measures, we estimated normalised Nash and Sutcliffe Efficiency (nNSE) which ranged between 0 and 1 providing more objective measure of model performance.

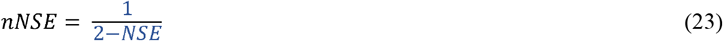

The *nNSE* accounts for the extent of variability in the observed values and provides more relevant values of model performance.

## Results and discussion

### Evaluation of model prediction

FAO-56 DCC was calibrated and validated for daily dynamics in the plant water availability (PAW) in the soil at all the study sites (48) over the three grapevine growing seasons (2018-19, 2019-20, 2020-21). A typical comparison of daily measured (M) PAW values in the soil profile estimated from moisture probe data and the corresponding FAO-56 DCC predicted values in relation to water content dynamics in the soil for site SR-A is shown in Fig. 3. While there were some differences in the observed and simulated values of water retained in the profile, but the moisture depletion pattern was almost similar across the season. At the time of bud burst in mid-September, PAW in the soil was close to the full capacity as a result of storage of winter rainfall during the dormant period. Water content in the soil gradually decreased thereafter in response to the water uptake by growing grapevines, evaporation losses, and downward movement of water in the soil. Irrigation was applied to provide adequate water supply to the grapevines to establish a desired canopy style during the bud burst to flowering (BB-FL) period.

**Fig. 3.**
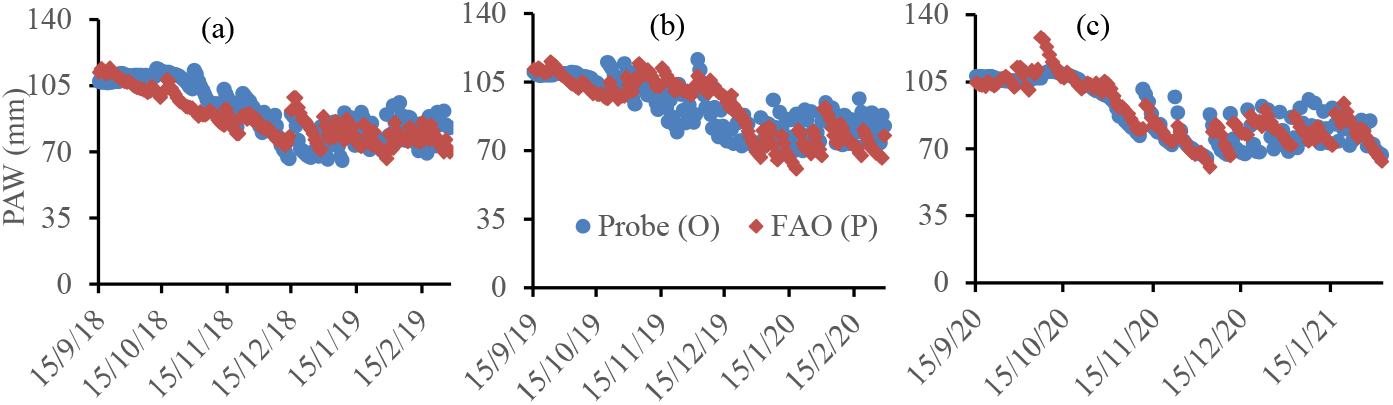
Comparison of daily measured (M) and predicted (P) real-time plant available water (PAW) in the profile for site SR-1A during the 2018-19, 2019-20 and 2020-21 grapevine growing seasons.

A gradual reduction in PAW indicated water deficit condition, especially during the December-January period which corresponded to applied stress over the FL-V growth stage to regulate the berry development and better biochemical composition which suits a characteristic wine quality (Castellarin et al., 2007; Cooley et al., 2017; Romero et al., 2019). Thereafter grapevine enter the berry ripening phase (V-H) during which a stable PAW in the soil was observed. Stable moisture regimes in the soil profile during V-H and post-harvest period coincides with appropriate level of stress applied with precise irrigation application for achieving the desired yield and yield attributes. This suggests that predicted water regimes in the soil profile corresponds well with the irrigation application, plant water uptake and stress applied during the entire grapevine season.

Statistical errors (RMSE, ME and MAE) were estimated to test the prediction of daily dynamics in water regime in the soil by the model at all the study sites during three consecutive seasons (2018-19, 2019-20, 2020-21) which is box-plotted in Fig. 4. The RMSE values across different sites ranged from 3.9-26.6 mm with a mean of 10.1 mm. These values remained below 10 % of the respective PAWC of the respective soils except site 14, 17 and 20 in both replicates. However, MAE and ME values showed relatively lower deviation as compared to RMSE and varied from 3-19 and -7.8 to 13.1 mm, respectively across all sites (24), replicates, and seasons (2018-19, 2019-20, 2020-21). It is worth mentioning that the sites showing inconsistent RMSE values showed better ME estimates. On the other hand, normalised Nash-Sutcliffe efficiency (nNSE) ranged from 0.3-1 showing a good simulation of daily PAW values in the profile by the model. Similarly, NSE and r^2^ values were also showed a good agreement between the measured and modelled water retention in the soil except couple of outliners and divergent values for sites 11A, 17A, 24A during 2018-19 season and at 14C during 2019-20 season. The average values of NSE, nNSE and r^2^ across sites and seasons were 0.68, 0.76 and 0.74, respectively showing a good agreement between measured and predicted values of PAW in the soil. This shows that the estimated water balance components by FAO-56 dual crop coefficient method match well with the measured data across all sites and seasons.

**Fig. 4.**
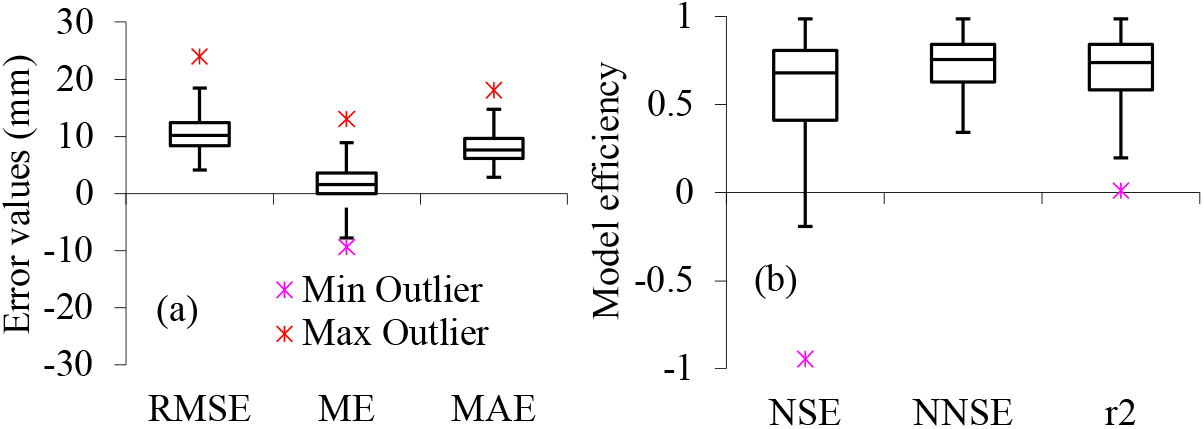
Error estimates (RMSE, MAE, ME), model efficiency (NSE), normalised NSE (nNSE) and coefficient of determination (r^2^) values between daily measured and predicted plant available water in the soil profiles under grapevines at different locations in the Barossa region.

FAO-56 DCC has been found to demonstrate similar prediction by several researchers in different crops comparing soil water contents (Fandiño et al., 2012; Paredes et al., 2017; Peddinti and Kambhammettu, 2019; Wu et al., 2016), *ET*_*c*_ estimated by eddy-covariance (Peddinti and Kambhammettu, 2019; Tian et al., 2016), and sap-flow measured actual transpiration *T*_*c act*_ ((Paço et al., 2011; Paço et al., 2019; Qiu et al., 2015). These studies show the reliability of FAO-56 DCC approach for estimating evapotranspiration components, crop coefficients and extent of stress imposed on the plant root water uptake under water deficit conditions. Some disparity in the measured and simulated PAW values was associated with insufficient prediction of deep drainage which was estimated as the excess amount of water over soil fill capacity.

### Seasonal irrigation and vine growth

There is a wide variability in the seasonal irrigation (*I*_*r*_) application (rainfed to 366 mm) across the Barossa zone (Fig. 5) depending on local variation in landforms, climate, genotype, soil characteristics, water availability and extent of stress imposed by the grower for establishing desired vine style, berry composition and distinct flavour, aroma, and quality of wine. For example, at SR11, irrigation application was more than 3 times of SR7 and SR13. Therefore, there was a lack of any temporal or seasonal trend in the irrigation data across the region, however seasonal average across study sites and seasons was 160 mm water applied through irrigation. Average seasonal rainfall during this period (15^th^ September to 31^st^ March) amounted to 114 mm. Numerous studies across the globe showed the impact of varied soil, rootstock, genotype and climate conditions on the seasonal irrigation applied to vineyards (Girona et al., 2005; Mancha et al., 2021; Munitz et al., 2019; Ohana-Levi et al., 2022; Ohana-Levi et al., 2020; Rallo et al., 2021; Romero et al., 2022).

**Fig. 5.**
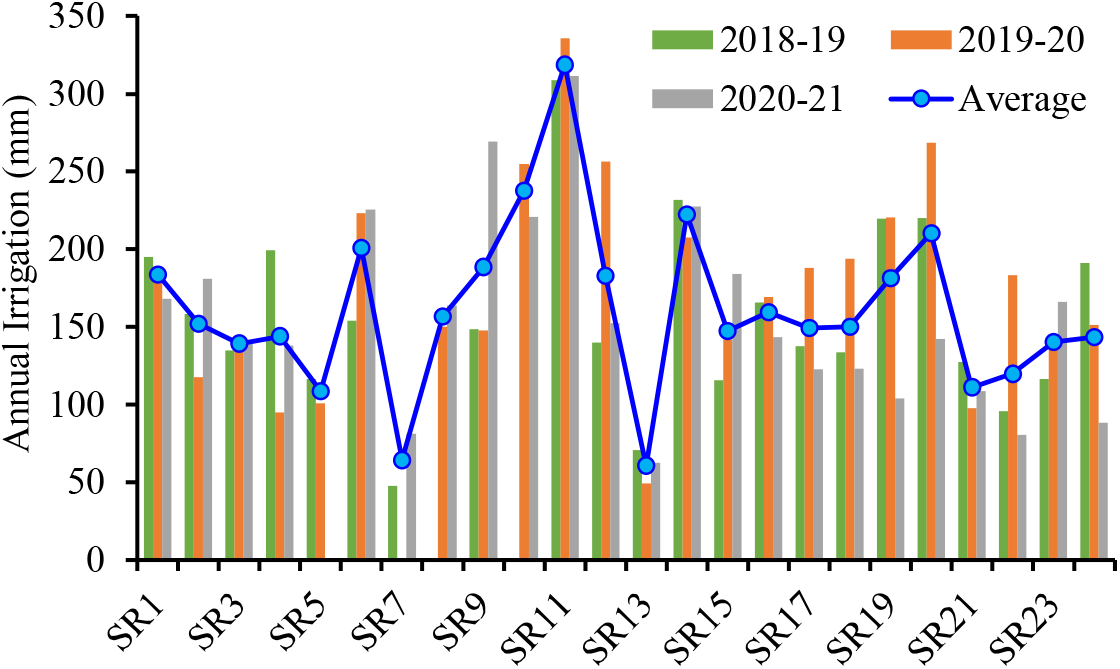
Irrigation amounts applied to grapevines at different sites (SR1 to SR24) during the 2018-19, 2019-20 and 2020-21 seasons in the Barossa and Eden Valley.

However, the average data showed an interesting revelation in the irrigation application over different crop growth stages. The mean irrigation amount across sites and seasons during the bud burst to flowering (BB-FL) period (15^th^ Sept - 15^th^ Nov) accounts for only 18.7 mm and ranged from 0-85 mm across the sites and seasons. Essentially, irrigation requirement during this phase is minimum as the soil profile contains enough moisture from the winter rain and adequately supplemented by rainfall during BB-FL period which ranged from 25.7-114 mm across different sites. Most of water requirement during the initial period contributes to the vigour development including shoots and leaves emergence and elongation which ultimately shapes the better canopy development.

Flowering to fruit set (FL-FS) is the most important stage for grapevine growth (16^th^ Nov – 15^th^ Dec). Water stress (pre-veraison) during this stage can have immense impact on flowering, fertilization and adequate fruit setting. Irrigation application ranged from 0-82 mm with a mean of 28.8 mm across the study sites representing 18% of the mean seasonal irrigation. Fruit setting is an inherent plant regulated process; however, a large fruit drop could occur if adequate water is not available in the crop root zone. Fruit set to berry development (FS-BD) phase requires considerable amount of water supply because an optimum berry development in size and numbers is essential yield trait of grapevines. Average irrigation application during this phase varied from 0-103.6 mm with a mean amount of 38.6 mm across the spatial and seasonal scale. It was interesting to note that average rainfall during this phase was only 8 mm across the study region. Water stress during this phase may hamper the growth and composition of the berries favourable for the formation of acids and phenols related to quality attributes and wine quality (Koundouras et al., 1999; van Leeuwen, 2008; Yang et al., 2022). The extent of impact on different cultivars depends on the severity and timing of water stress (Alexander et al., 2020).

Meanwhile, the berry development to maturation known as veraison (BD-V) period (8^th^ Jan-25^th^ Jan) play a key role in the further development of quality traits in the berries. However, the berry development including Veraison phase spread over the entire period starting from fruit-set to fruit maturation which is a characteristic feature of different genotypes. This phase requires moderate level of water stress depending on the genotype being grown and excessive amount of irrigation application may degrade the quality attributes of mature berries. Irrigation applied to grapevines, mostly Shiraz in the study region, during this phase varied from 0-105 mm with a mean value of 36.2 mm over the 3 consecutive seasons (2018-19 to 2020-21). On the other hand, veraison to harvest (V-H) phase received 37.9 mm average irrigation which was higher than the veraison phase indicating some inconsistencies in the irrigation application during this period (Junquera et al., 2012).

Numerous studies across the world have made considerable efforts to understand and optimize the irrigation requirement of grapevine and extent of water stress applied during the different growth stages for realizing better wine quality. Different deficit irrigation strategies such as sustained (SDI) and regulated (RDI) deficit and partial root zone drying (PRD) have been evolved and applied to improve the water use efficiency, vine vigour, berry composition and wine quality including the production of healthier wines with lower alcohol content and higher concentrations of anthocyanins, tannins and other potentially beneficial organic compounds (Chaves et al., 2010; Ferrandino and Lovisolo, 2014; Intrigliolo and Castel, 2009; Ju et al., 2019; Markus et al., 2016; McCarthy, 2000; Pérez-Álvarez et al., 2021; Romero et al., 2019; Romero et al., 2022; Yang et al., 2022).

### Seasonal evapotranspiration, transpiration, and evaporation

Model predicted average daily values of actual evapotranspiration (*ET*_*C act*_), actual transpiration (*T*_*p act*_) and evaporation from exposed soil surface (*E*_*s*_) and respective deviation across study sites and seasons (2018-19 to 2020-21) are shown in Fig. 6. The overall average seasonal *ET*_*C act*_ for the study region accounts for 325.2 mm of water use with 30.7, 17.8, 16.3, 13.4, and 21.8 % of ET losses occurring during the BB-FL, FL-FS, FS-BD, BD-V, V-H period of grapevine growth, respectively. However, the seasonal *ET*_*c act*_ varied from 119-546 mm across different sites and seasons. Variability among the *ET*_*c act*_ values represents the variable water regimes, crop management and variable water stress imposed to obtain a good berry composition which differs at different locations. Similar seasonal *ET*_*C act*_ losses were observed for wine grape by Phogat et al. (2017) in the same region and by (Cancela et al., 2015) for Mediterranean climate, different soil and different vine genotype and style. (Yunusa et al., 1997) reported ET similar to current study for own-rooted and grafted Sultana grapevine under Australian conditions. However, (Williams et al., 2010a) observed much higher *ET*_*C*_ (574-829 mm) for lysimeter-planted Thompson seedless grape vine and more than double the amount of irrigation applied compared to our study.

**Fig. 6.**
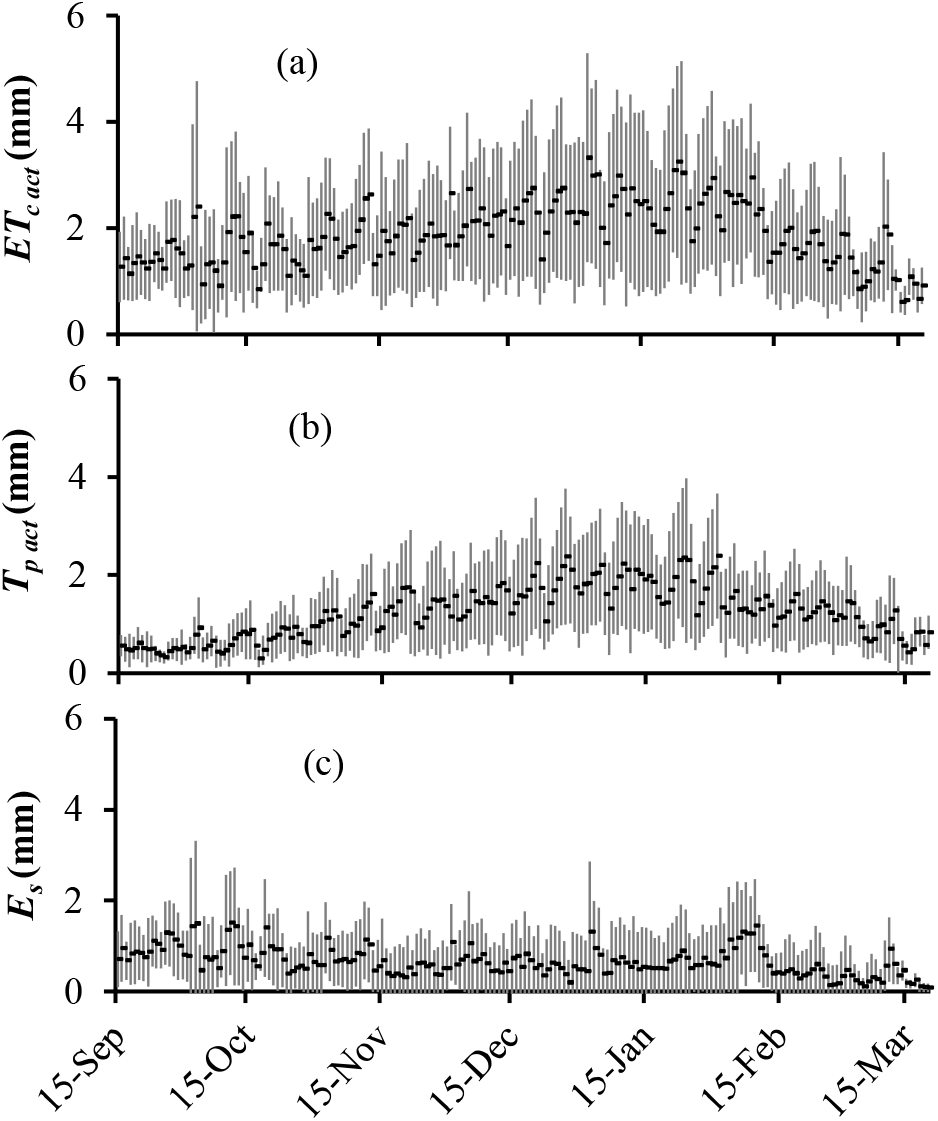
Predicted average values and the deviation in the actual evapotranspiration (*ET*_*C act*_), transpiration (*T*_*p act*_) and evaporation (*E*_*s*_) components of grapevines across study sites and seasons (2018-19 to 2020-21) in the Barossa region. Dots represent average values and vertical bars show the standard deviation based on year-to-year variability.

The daily average *ET*_*C act*_ values varied from 0.49-3.21 mm/day with a 20-97% variability across the region in the respective daily average. The maximum average daily value of 5.2 mm/day was obtained during mid-January in a high vigour grapevine irrigated frequently. Average grape vine daily *ET* rate from nine international field experiments was 3 mm/day (Teixeira et al., 2007), which represents varied rootstocks and mostly surface irrigation conditions. Other studies (e.g., (Fandiño et al., 2012; Shapland et al., 2012; Yunusa et al., 1997) showed average daily *ET*_*C*_ varying between 2-2.2 mm/day. Average monthly *ET*_*c act*_ for November, December and January months was 21, 22 and 28% (Fig. 7) of the seasonal value totalling more than 60% water use during this period conceding the maximum crop requirement because of hot and dry weather conditions.

**Fig. 7.**
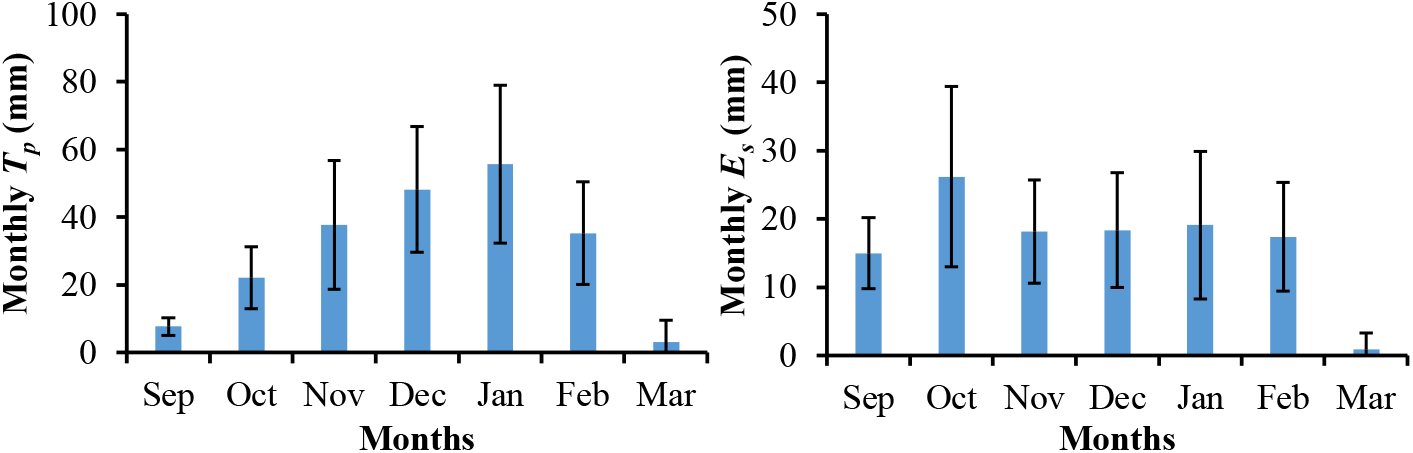
Monthly average transpiration (*T*_*p act*_) and evaporation (*E*_*s*_) under wine grape and their respective deviation across the study sites in the Barossa region.

Similarly, the overall average seasonal *T*_*p act*_ that accounts for 65% water losses in the seasonal *ET*_*C act*_ of grapevine across different sites and seasons (Fig. 6b, c). Average transpiration across different sites varied from 47-396 mm with 54% water uptake during FL-V period. However, wide variability was observed in the daily and seasonal values at different locations and seasons. The average daily values varied from 0.3-2.3 mm/day. Monthly average *T*_*p act*_ for December and January months accounted for 48.2 and 55.7 mm, transpiration losses respectively (Fig. 7) representing 19 and 23% of total seasonal water use by the grapevine. Ratio of actual transition and the evapotranspiration losses (T_c act_/ET_c act_) is highly variable and depends on numerous factors such as seasonal climate, genotype, soil type and water stress imposed by the growers leading to a highly variable canopy cover and evaporative flux from the soil surface. However numerous studies have reported similar extent of ET_c act_ and T_c act_ from grapevines (Sánchez et al., 2019).

Average seasonal *E*_*s*_ accounts for 35% (115 mm) of the total *ET*_*c act*_ and 46% of that occur during the BB-FL period of grapevine growth ranged from 25-68% across study sites depending on the soil surface conditions (Fig. 6). During this period a large, wetted surface area is exposed to atmosphere contribute to large amount of unproductive water directly escape to the atmosphere. Daily *E*_*s*_ losses ranged from 0-5.9 mm/day across different sites. These losses consequently decreased as the canopy cover increase and wetted area near the tree line exposed to the atmosphere decreased. About half of the seasonal *E*_*s*_ (46%) is occurring during the BB-FL stage of vineyard growth and another 15% and 20% respectively during FL-FS and V-H stage of growth. Higher evaporative losses during the full canopy stage are minimum which has been attributed to full grown canopy.

Numerous studies have reported similar or higher evaporation losses under the drip irrigated grapevines. (Valentín et al., 2023) reported that 21-50 % of *ET*_*c act*_ corresponds to seasonal *E*_*s*_ losses.

### Single, basal and evaporation coefficients

Average daily values and corresponding deviations in climate adjusted (*K*_*c adj*_) and actual single (*K*_*c act*_) crop coefficients are compared with the standard generic Kc curve for grapevines in Fig. 8a. It should be noted that standard values (*K*_*c std*_; Allen et al., 2009) have been adjusted for the corresponding location specific climate variability and soil condition (*K*_*c adj*_) which were further adjusted for real-time grapevine (*K*_*c act*_) growth at different locations. Interestingly, daily *K*_*c adj*_ values during mid and late season were distributed in close proximity to the corresponding *K*_*c std*_ values. The average *K*_*c adj*_ values for mid (0.69) and late (0.53) grapevine season across different sites and seasons were similar to *K*_*c std*_ values. However, during the initial growth stage, there was a wide variability in the *K*_*c std*_ and *K*_*c adj*_ values due to a varied existing surface conditions at different locations in the study region.

**Fig. 8.**
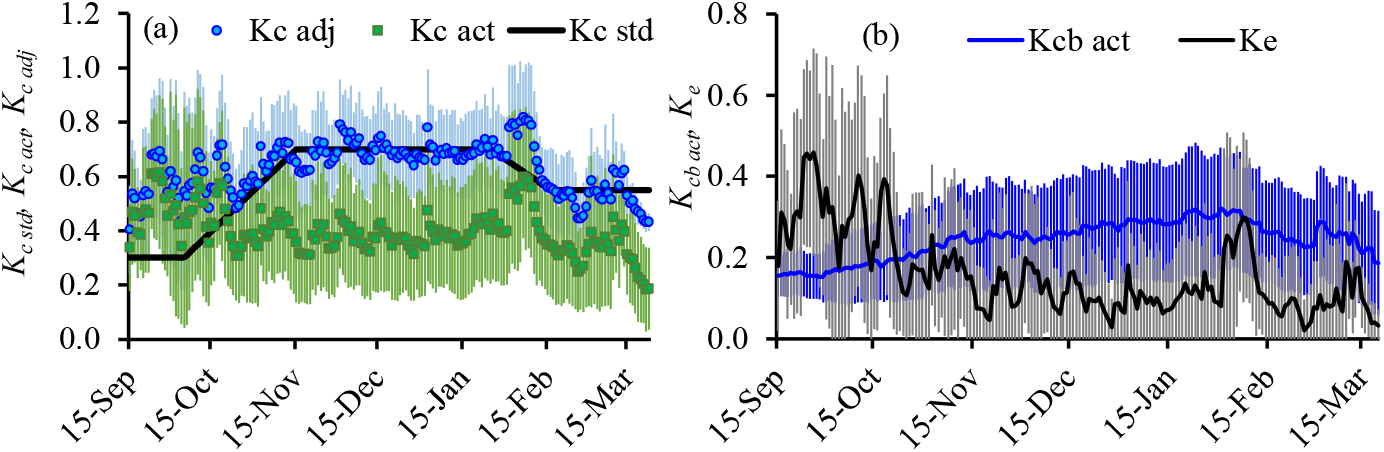
Spatial variability in single crop coefficient (*K*_*c act*_), basal crop coefficient (*K*_*cb act*_) and evaporation coefficient (*K*_*e*_) of grapevines in the Barossa region. Solid lines represent the mean values and vertical bars represent the standard deviation across mean based on year-to-year variability.

Actual single crop coefficient (*K*_*c act*_) values reduced drastically in response to local climate, soil and exposure to seasonal water stress (Fig. 8a). During the BB-FL stage of crop growth, average values of *K*_*c act*_ across all sites varied from 0.35 to 0.59 showing a variation of 3 to 53% (mean CV 29.8%) among the average values at different sites. Similarly, during the FL-V stage of growth the average values of *K*_*c act*_ across the sites varied from 0.16-0.62 with CV values ranging from 7 to 43% with mean variation of 19.6% indicating less variability among the *K*_*c act*_ across the region. On the other hand, the average *K*_*c act*_ during the V-H stage of grapevine growth ranged from 0.18-0.68 indicating slightly higher ET flux across the region than during FL-V period. The variation among the *K*_*c act*_ values during this phase ranged from 10-56% with a mean deviation of 26.7% across the sites. This is similar to the variability observed during the initial phase but higher than the mean value during the FL-V stage of crop growth. The *K*_*c act*_ values represents the sum of respective *K*_*cb act*_ and *K*_*e*_ estimated by the FAO-56 DCC approach. It should be noted that *K*_*cb act*_ curve stabilised across the study sites with CV varied from 4.6-26.7% with a mean deviation of 17% (Fig. 8b). The estimated average *K*_*cb act*_ values for the initial, mid and late stage of grapevine growth representative of the Barossa region in accordance with FAO-56 (Allen et aal., 1998) were 0.45, 0.38 and 0.37, respectively. However, the daily variation in the *K*_*cb act*_ across 48 study sites over the 3 cropping seasons contains enormous variability especially during the early stage of crop growth (BB-FL) which is assimilated in the respective *K*_*c act*_ values too.

It is interesting to note that average *K*_*e*_ values across all the sites showed almost 50% evaporation contributed during the initial growth stage of grapevine in the Barossa zone. The average *K*_*e*_ values varied from 0.18-0.38 (0.25), 0.06-0.16 (0.11), 0.07-0.25 (0.14) with CV ranging from 15.7-61.1 (30.8%), 7.6-63.9 (30.6%), and 8.4-51.6 (34.5%) %, respectively during the BB-FL, FL-V and V-H stages of crop growth (Fig. 8b). The average values during the FL-V and V-H stages are similar to the Es losses (0.15) reported FAO-56 (Allen et al., 1998). Similar deviation in the *K*_*e*_ values were reported by (Phogat et al., 2020a) in the area close to the current study region under historic and future climate projections. Other studies (Cancela et al., 2015; Poblete-Echeverría and Ortega-Farias, 2013) also reported similar location specific estimation of evaporative losses. It suggests that the evaporation component of the evaporative flux is highly variable as compared the water losses through the transpiration stream. This is probably related to much more variation in the water dynamics at the soil surface than within the soil profile. During this phase canopy is not fully developed leaving a large, wetted area of the soil surface exposed to direct atmospheric losses. Early stage of crop with low canopy cover encourages higher wetted surface area leading to higher water losses. This is the probably the most crucial stage of growth requiring appropriate control measures to reduce the unwanted water losses.

Daily values of *K*_*c act*_, *K*_*cb act*_ and *K*_*e*_ averaged across the study sites and seasons is shown in Fig. 9. Overall, the variability in the *K*_*c act*_ values was higher during the initial period of growth which is mostly contributed by the *K*_*e*_ component. During this stage higher wetted surface is exposed to the water losses to the atmosphere as the canopy cover is at its minimum. Similarly high evaporation during the first half of February also contributed by the higher *K*_*e*_ component. *K*_*cb act*_ curve drawn from the daily average values across sites and seasons depicts a smooth transition during the season.

**Fig. 9.**
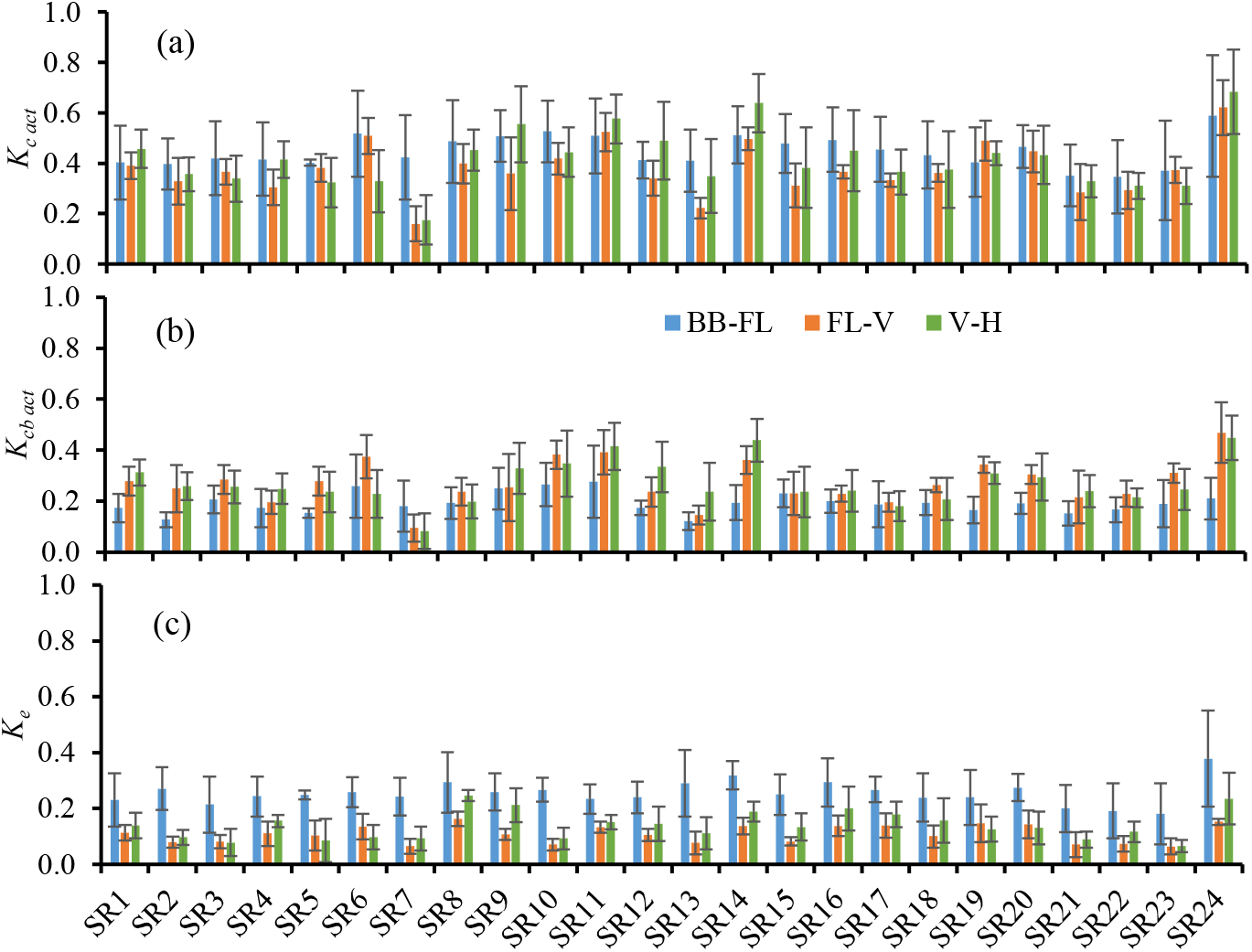
Model predicted average values of (a) actual single crop coefficients (*K*_*c act*_), (b) basal crop coefficients (*K*_*cb act*_) and (c) evaporation coefficients (*K*_*e*_) during bud-burst to flowering (BB-FL), flowering to Veraison (FL-V) and Veraison to harvest (V-H) stages of wine grape growth at different sites (SR1 to SR24) in the Barossa region.

Among the sites, SR6, 9-11, 14 and SR24 showed relatively higher K_c act_ values while SR7, 13, 21-23 showed relatively lower values. It was observed that the former sites received relatively higher amount of irrigation as compared to the latter group. Maximum values of *K*_*c act*_ during BB-FL, FL-V and V-H stages were observed at SR24, 24, 24 and lower values at SR21, 7 and 7, respectively. Almost similar trend was observed in the *K*_*cb act*_ values during the various stages of growth (bud burst to harvest) across different study sites. Average *K*_*cb act*_ values during BB-FL, FL-V and V-H varied from 0.12-0.28, 0.10-0.47 and 0.08-0.45 with CV varying from 12-55 (32.8%), 9-57(24.4%) and 13.7-85% (31.4%).

### Vineyard water stress dynamics

Water stress coefficient (*K*_*s*_) is a dimensionless transpiration reduction factor dependent on the availability of water in the soil profile. The average stress coefficient (*K*_*s*_) values during the BB-FL period (late autumn to early summer period) varied from 0.44-0.69, lower values mean higher stress (fig. 10). Normally the soil contains lot of water during this phase as this period overlaps post rainy season in the Mediterranean climate. The extent of water availability depends on the rainfall received during the dormant period and water holding capacity of soils. The average level of water stress increased by 20% during the FL-V phase and varied from 0.17-0.74 across different study sites. Similarly, the stress level during the V-H period of growth reduced by 15% and varied from 0.17-0.83. Values of *K*_*s*_ at SR-7 site were the lowest indicating very high-water stress over the growing season because the irrigation application was the lowest among the sites (50-80 mm).

**Fig. 10.**
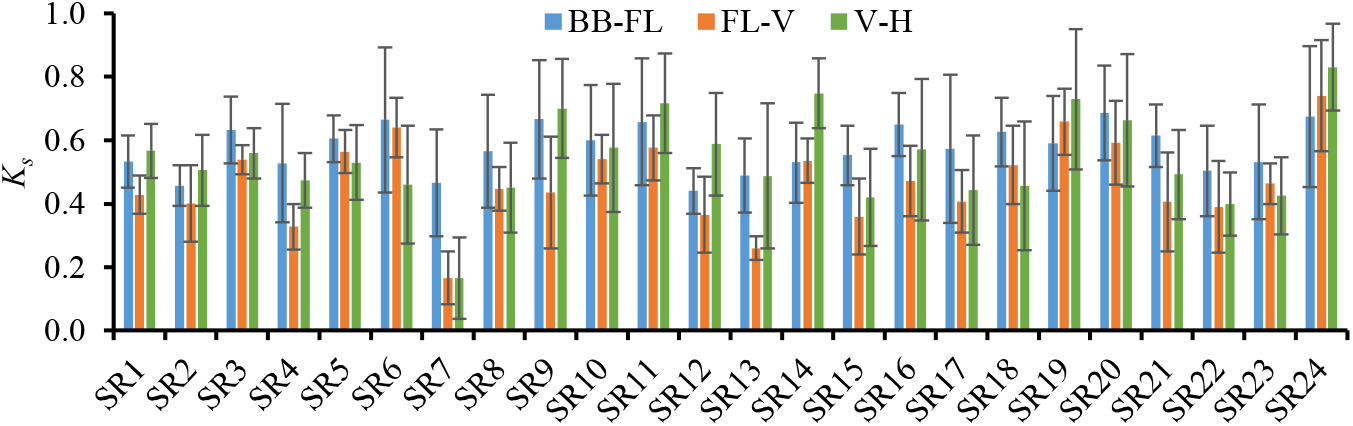
Average water stress coefficient (*K*_*s*_) values at different study sites (SR1-SR24) during the bud burst-flowering (BB-FL), flowering-veraison (FL-V) and veraison-harvest (V-H) periods of grapevine growth.

The relative stress level and the shape of the *K*_*s*_ curve determines the magnitude of the effect of the stress on the process between the thresholds. The relative stress is 0.0 at the upper threshold and 1.0 at the lower threshold (Figure 10). The shape of most of the *K*_*s*_ curves are typically convex as shown in the Fig. 11. Initially the *K*_*s*_ fluctuates around 0.7 which is normal threshold outlined in FAO-56 (Allen et al., 1998) for grapevines. The *K*_*s*_ gradually decreased as the season progressed and attained the lowest average value between Mid-November to Mid-December which corresponds to berry development phase. Water stress during this period can have drastic impact on the growth and composition of berries.

**Fig. 11.**
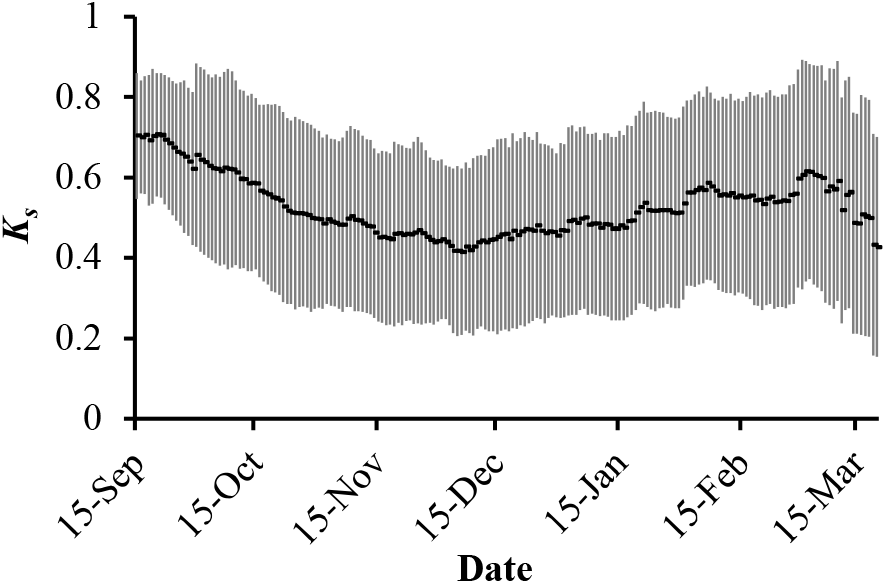
Variability in the stress coefficient (*K*_*s*_) imposed to the irrigated grapevines in the Barossa region. Dots represent the mean values and vertical bars represent the standard deviation across mean based on spatial heterogeneity at regional scale.

Extent and timing of water application play a crucial role in growth and yield as well as wine quality. Primary focus of the Barossa region is on producing high quality wine requiring depends on the extent of pre- and post-veraison stress applied. Extent of pre-veraison stress could cause more post-veraison stress due to the cumulative stress. However, sustained water deficit conditions (similar stress during pre- and post-veraison) over the season have as cumulative seasonal impact including reduced canopy growth, yield and grape quality (Girona et al., 2009; Wenter et al., 2018).

Mild or moderate water deficits at this stage generally favour the berry accumulation of sugar and some phenolics (e.g. anthocyanin), whereas severe water stresses can lead to significantly reduced berry quality (sugar, aroma) and grape yield (Van Leeuwen and Destrac-Irvine, 2017; van Leeuwen, 2008; Yang et al., 2022). The effect of water availability varies with the variety. Limited water supply can increase skin phenolics (e.g. anthocyanins) which is crucial for the production of high-quality red grapes, whereas this may negatively affect the aromas of white grapes (van Leeuwen, 2022; van Leeuwen and de Rességuier, 2018). However, the effects of water deficit also depend on its timing and duration (Bonada et al., 2021; Gambetta et al., 2020). Before veraison, berries are hydraulically connected to vine and are sensitive to drought-induced shrivelling, but they are largely insensitive at/after veraison since berries become buffered from changes in vine′s water status (Gambetta et al., 2020).

Water shortage before veraison can have strong negative impacts on leaf growth, berry weight and final yield per vine (Chacón-Vozmediano et al., 2020; van Leeuwen, 2022; Wenter et al., 2018; Yang et al., 2022). One of the most important pre-veraison phenophase corresponds to the flowering-veraison phase, which proves to be an important period for berry weight and yield determinations (Chaćon (Chacón-Vozmediano et al., 2020; Yang et al., 2022). For instance, (Ramos and Martínez-Casasnovas, 2014) found grape yield to be particularly sensitive to water availability during bloom-veraison period while (Yang et al., 2022) reported that an increased potential yield loss rate during this phase corresponds to variety characterization, soil and climate conditions, however, the impact of water stress depends on its timing and duration (Gambetta et al., 2020). It is well understood that water stress before veraison can have strong negative impacts on leaf growth, berry weight and final yield per vine (Chaćon-Vozmediano et al., 2020; Gambetta et al., 2020; van Leeuven et al., 2018; Wenter et al., 2018) especially on berry weight and yield determinations (Chaćon-Vozmediano et al., 2020; Gambetta et al., 2020; Ramos and Martinez-Casasnovas, 2014). Therefore, it is important to focus on how to increase water availability and mitigate possible water stress over this period, while not using irrigation when it is restricted.

### Water productivity and yield-water stress relationship

Water productivity (WP) is an important parameter which shows the extent of biomass (b) and/or berry yield (y) produced with a unit expense of water. Extent of water usually assumed as the actual crop evapotranspiration (*ET*_*C act*_) which also includes the unproductive water lost from the soil surface (*E*_*s*_). For more precise estimation of WP, water lost through the plant transpiration stream (*T*_*p act*_) is considered as the actual expanse of water consumed to produce certain amount of photosynthates. Estimated average water productivity for biomass (WP_ET_C_b) and berry yield (WP_ET_C_y) in relation to ET_C_ ranged from 0.78-5.36 and 0.18-1.28 kg/m^3^/ha, respectively (Fig. 12). Similarly, the estimated average WP for biomass (WP_T_p_b) and yield (WP_T_p_y) in relation to actual transpiration (T_p act_) was higher than respective Wp_ET_C_ at all the sites and ranged from 1.17-8.37 and 0.37-2.0 kg/m^3^/ha, respectively. Among the sub-regions, SG (sites 13-16) and EE (sites 9-12) showed higher water productivities (biomass and berry yield) than the average WP across the Barossa region. In SG, WP_ET_C_b and WP_ET_C_y was 55 and 38% higher than the regional average of 0.54 and 2.28 kg/m^3^/ha, respectively (Fig. 12). High production water productivity in SG is related to more water availability to the grapevines in the soil as Southern Grounds had the highest plant available water capacity (PAWC) in the soils which provided continuous water supply for higher yields as compared to other sites (Phogat et al., 2024). Water productivity for biomass and yield in reference to vine transpiration losses also showed similar response. On the other hand, average WP for CG sub-region (sites 5-8) was lower than the corresponding average values of the entire Barossa region. Average regional values of WP_ET_C_b, WP_ET_C_y, (WP_T_p_b), and (WP_T_p_y) were 0.54, 2.28, 0.85 and 3.59 kg/m^3^/ha, respectively. Average water productivity for biomass as well as berry yield relative to transpiration increased in a similar proportion (around 57-58%) across different sub-regions compared to WPETc indicating a similar response of evaporation losses across different sites.

**Fig. 12.**
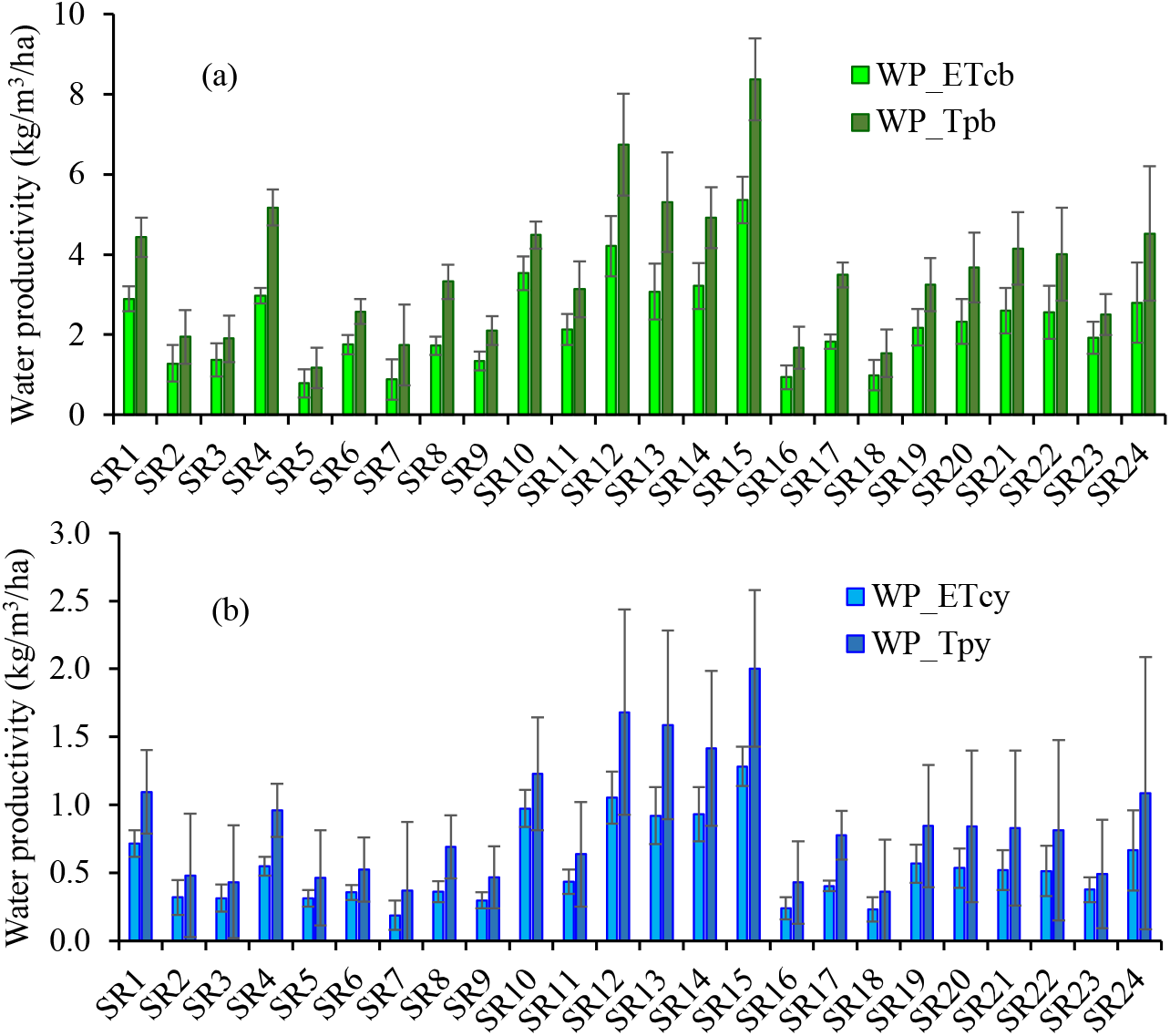
Estimated water productivity in relation to actual evapotranspiration (WP_ETc) and actual transpiration (WP_Tp) for (a) biomass and (b) berry yields at different sites.

Several studies reported a varied extent of water use efficiency by grapevines. Stevens et al. (2010) reported that reduced irrigation did not affect the irrigation water use index, whereas other investigations (Fereres and Soriano, 2006; Williams et al., 2010b) reported that deficit irrigation does increase water use efficiency for woody perennial crops. However, these studies used different measures of water use for estimating water use efficiency. In contrast, Phogat (Phogat et al., 2017) illustrated a clear relationship between a reduced water application and increased water productivity. They found that application of 50% sustained deficit irrigation increased the water productivity by 21% as compared to well-watered grapevines and water productivity relative to transpiration (*WPT*_*p*_) was approximately double than *WPET*_*C*_, which indicates high Es losses in this study. Comparable grapevine water productivities (1.8 kg/m^3^) against *ET*_*C*_ were recorded in other studies for similar amounts of water (Atroosh et al., 2013). While very high values of water productivities were reported in other studies (e.g., (Phogat et al., 2017; Williams et al., 2010b) under SDI irrigation of grapevines.

## Conclusions

Accurate estimation of crop water requirement is crucial for enhancing water productivity, yield and quality attributes as well as understanding the temporal and seasonal trends in crop water needs. Locally estimated crop coefficients and water balance components serve as reliable parameters for estimating accurate water needs of irrigated corps. Results showed a wide variability in the crop coefficients values due to a varied climate and soil conditions at different locations in the study region. Estimated average actual crop coefficients (*K*_*c act*_) across all sites varied from 0.35 to 0.59, 0.16-0.62 and 0.18-0.68 showing a variation of 3 to 53, 7-43, and 10-56%, respectively across the study region. High variability in the crop coefficients during the growth stages induced a lack of any temporal or seasonal trend in the irrigation requirement for grapevines across the region, however seasonal average across study sites and seasons was 160.2 mm. Nevertheless, crop coefficients and stress coefficients estimated for different sites can be used to schedule irrigation and to regulate the timing of water stress applied in the grapevines across the Barossa region.

Results further showed higher pre-veraison water stress across the region which may induce a cumulative seasonal impact including reduced canopy growth and yield for producing high quality wines. It is well understood that the effects of water deficit on grapevines depend on its timing, extent and duration of exposure to stress. Before veraison, berries are hydraulically connected to vine and are sensitive to drought-induced shrivelling, but they are largely insensitive at/after veraison since berries become buffered from changes in vine′s water status.

FAO-56 dual crop coefficient approach has been able to simulate the real-time soil moisture regime in soils influenced by irrigation, crop water use and extent of water stress applied over the multiple grapevine seasons.

Overall, the results from this study demonstrated that predicted variability in actual evapotranspiration, transpiration and evaporation not only corresponds to climate and soil water regimes but also highlights the key management decisions implemented by the individual grower for achieving the better-quality traits (desired vine style, berry composition and distinct flavour, aroma, and quality of wine) rather than achieving higher yield. Thus, varied extent of inter- and intra-season water stress (pre-veriason and post-veriason) at different locations imposed by the managers imply that these decisions are more of site-specific rather than imposed uniformly across the irrigated sub-regions and region.

## Acknowledgements

We thank Mr. Gaston Sepulveda, Mr. Lucas De Simoni, and Mrs. Annette James for their support in the collection of soil cores and the broader research team for their support. We are grateful to the growers of the Barossa region for providing access to their vineyards. This work was supported by Wine Australia, with levies from Australia’s grape growers and winemakers and matching funds from the Australian Government. The South Australian Research and Development Institute and the University of Adelaide are the members of the Wine Innovation Cluster in Adelaide.

## Notes

### Competing Interest Statement

The authors have declared no competing interest.

### Summary of Updates

This version is revised due to the discrepancy in the previous version.

